# Under conditions of high wall shear stress, several PfEBA and PfRH ligands are important for *Plasmodium falciparum* malaria blood-stage growth

**DOI:** 10.1101/2024.09.24.614720

**Authors:** Emma Kals, Morten Kals, Pietro Cicuta, Julian C. Rayner

## Abstract

Malaria kills over 600,000 people annually, with all the clinical symptoms being caused by the blood stage of the infection. Malaria parasites invade red blood cells (RBCs), where they grow and multiply until daughter parasites egress to invade new RBCs. This cycle happens primarily in the blood circulation, bone marrow, and spleen, where parasites and RBCs are exposed to flow-generated forces. Despite this, almost all *in vitro* growth assays are carried out in static conditions, which are a poor mimic for the conditions that malaria parasites encounter in the body. Therefore, we systematically tested the impact of dynamic conditions created by orbital shaking platforms on parasite growth and explored the link between growth and the wall shear stress forces generated by fluid motion. For several strains of the deadliest malaria species, *Plasmodium falciparum*, we showed that strikingly, for any given vessel, there is a critical shaking speed at which growth rates are reduced, corresponding to when the RBCs start to aggregate in the centre of the well, before growth increases again at higher shaking speeds. The force that parasites are exposed to at this critical shaking speed corresponds cloesly with previous measurements for the forces that RBCs and parasites are exposed to in the microvasculature. During invasion, the early attachment of the parasites to RBCs is dependent on two families of attachment proteins, the Erythrocyte Binding Antigen (PfEBA) family and the Reticulocyte Binding Protein Homologue (PfRH) family, which are thought to be largely redundant in function. We used a panel of PfEBA and PfRH knock-out lines to show for the first time that several of these ligands have greater importance in high wall shear stress conditions. This both adds new understanding to the function of these ligand families, and indicates that the concept of the critical shaking speed can reveal new parasite growth phenotypes.

## 1 Introduction

All clinical symptoms of *Plasmodium falciparum* malaria are caused by the blood stage of the infection, during which the parasites invade and replicate inside red blood cells (RBCs). Blood stage growth requires parasites inside mature infected RBCs, known as schizonts, to egress, releasing the merozoite form of the parasite that goes on to invade uninfected RBCs, a process that typically takes less than a minute [1]. After invasion, the parasite develops into a new schizont containing 26-32 individual merozoites over the next 48 hours [2, 3]. Where parasites develop and invasion occurs in the body is not completely understood; post-mortem analysis of patients with severe malaria has shown that there is a high attachment due to cytoadherence of infected RBCs to capillaries and postcapillary venules in organs [4], and so a key site of invasion is likely in the subsequent vasculature. There is also increasing evidence that a significant proportion of parasite burden may be in the bone marrow [5] and spleen [6, 7]. In these different environments, both parasites and RBCs will be exposed to shear forces resulting from gradients in the velocity, intrinsic to fluid flow in confinement, but *in vitro* studies of parasite growth and invasion have primarily been performed only under static conditions, which is an imperfect mimic of the *in vivo* environment.

Shaking on an orbital rotation platform is a simple and low-cost method of introducing motion in parasite cultures. While it does not accurately mimic physiological conditions, orbital shaking creates shear forces comparable to those experienced in the vasculature and is commonly used when culturing epithelial cells [8]. The fluid motion induced by shaking can be characterized by two fundamental flow regimes: laminar and turbulent. In laminar flow, fluid moves in smooth, orderly layers, creating shear forces that primarily arise from the interaction between the fluid and the container walls, leading to a gradual velocity gradient. In contrast, turbulent flow is characterized by chaotic, irregular motion with significant mixing between fluid layers, resulting in more complicated shear forces. Both flow types exert shear stress on the flask walls, referred to as wall shear stress (WSS), which will influence the interactions between parasites and red blood cells (RBCs) in sedimented culture.

Several studies have reported that shaking influences the growth and behaviour of *P. falciparum* in culture, though results have been inconsistent, likely due to varying methodologies and unquantified forces. Shaking has been shown to either enhance [9, 10, 11, 12, 13] or reduce growth rates [14]. Shaking also distributes parasites more evenly through the culture, reducing the number of RBCs infected by multiple parasites [15, 14, 16, 12]. However, the absence of systematic comparisons across a range of shaking speeds and flow regimes makes it challenging to untangle the precise effects of shear forces on parasite growth.

Vessel shape and size, fluid volume, fluid viscosity, orbital radius, and orbital rotation speed will all impact fluid motion and the resultant mechanical forces generated, and without these being clearly described, it is impossible to draw clear conclusions from the existing literature. It is also worth noting that flow conditions at a given speed will vary across a well due to the non-uniform shear stress field generated by the orbital motion [17], which means complex models are needed even to predict the WSS exerted on the base of the wells [18, 19, 20, 8]. By systematically testing a range of different conditions at different orbital shaking speeds, we show that growth under shaking is strongly impacted by shaking speed, vessel geometry and haematocrit. This allowed the systematic exploration of the effects of shaking on parasite growth in both laminar and turbulent conditions. This established that shaking can both decrease and increase growth and identified a critical shaking speed at which the blood starts to move into suspension. At critical shaking speed, the magnitude of the WSS is comparable to that of the microvasculature. This makes the comparison of growth in static and at the critical shaking speed an interesting and novel phenotyping assay for understanding the role of different invasion ligands.

During malaria parasite invasion of RBCs, two families of proteins are thought to be involved in the initial attachment of the parasites, called merozoites, to RBCs, the Erythrocyte Binding Antigen (PfEBA) family and Reticulocyte Binding Protein Homologue (PfRH) family of ligands. None of the proteins are essential as deleting them individually in the *P. falciparum* genome alone does not block invasion [21, 22] with the exception of PfRH5 that has a distinct and essential role at a later step int he invasion process [23, 24]. It is widely hypothesised that PfEBAs and PfRHs are functionally interchangeable [25, 26], and that variation in expression of PfEBA and PfRH proteins allows the parasite to utilise different receptors and therefore different invasion pathways. This is perhaps to help evade the host immune response and/or allow parasites to overcome variation in surface receptors [27, 28, 29]. However, the range of phenotype approaches that PfEBA and PfRH knockout lines have been exposed to has been relatively limited, primarily involving either flow-cytometry or video microscopy-based invasion assays carried out in static growth conditions. We therefore applied the concept of the critical shaking speed to a library of genetically modified *P. falciparum* lines in which PfEBA and PfRH ligands have been deleted, and established that several of these ligands are more important when the invasion happens under high-shear stress conditions. This adds to our understanding of these critical gene families, and suggests that another reason for the large number of different PfEBA and PfRH ligands is to allow invasion under different shear stress conditions in different regions of the circulatory system.

## 2 Methods

### 2.1 *Plasmodium falciparum* culture

*P. falciparum* strains (either the wildtype NF54, 3D7 or Dd2 lines, or genetically modified lines) were cultured in human erythrocytes purchased from NHS Blood and Transplant, Cambridge, UK – ethical approval from NHS Cambridge South Research Ethics Committee, 20/EE/0100, and the University of Cambridge Human Biology Research Ethics Committee, HBREC.2019.40, formal written consent was obtained by the NHS. Unless indicated cultures were kept at a 4% haematocrit (HCT). Cultures were maintained in RPMI 1640 media (Gibco, UK) supplemented with 5 gl^−1^ Albumax II, DEXTROSE ANHYDROUS EP 2 gl^−1^, HEPES 5.96 gl^−1^, sodium bicarbonate EP 0.3 gl^−1^ and 0.05 gl^−1^ hypoxanthine dissolved in 2M NaOH. Culture was performed at 37 ° C in a gassed incubator or gassed sealed culture container under a low oxygen atmosphere of 1% O_2_, 3% CO_2_, and 96% N_2_ (BOC, Guildford, UK).

### 2.2 Sorbitol synchronisation

Sorbitol synchronisation was performed as described by [30]. Ring stage cultures were pelleted by centrifugation, the supernatant was removed, and the pellet was resuspended at ten times the pelleted volume of 5% D-sorbitol (Sigma-Aldrich), then incubated at 37 ° C for 5 mins, during which time later stage parasites (trophozoites and schizonts) rupture. The cells were pelleted by centrifugation and then resuspended in RPMI at twenty times the pelleted volume. The cells were pelleted again and then resuspended in complete media to give 4% HCT.

### 2.3 Preparation of synchronised parasites for growth assay

All lines were cultured in static conditions prior to the assay. For each run of the assay all lines were defrosted at the same time, sorbitol synchronised in the first cycle when the culture was greater than 0.5% rings and then delayed at room temperature to push the egress window to the intended start window. Smears were checked to confirm the culture was predominantly schizonts with few rings. All but 5 ml of the media was removed, and the infected blood was resuspended in the residual liquid, then layered over 5 ml of 70% Percoll in a 15 ml tube (Percoll Merck P4937, 10% 10x PBS, 20% RPMI) and centrifuged at 1450 rcf for 11 min with break set to 1 and accelerator to 3. The band of late-stage parasites at the Percoll-media interface was then added to a flask with media and blood at a 4% HCT and incubated in a gassed incubator at 37 ° C for 3 h. The Percoll separation was then repeated as above, but this time, the bottom pellet containing ring stage parasites (which had invaded in the previous 3 h) was kept, and then sorbitol synchronised to completely eliminate any late stage parasites, resulting in tightly synchronised parasites within a 3h development window. The parasitemia was then counted using Giemsa smears and the cultures diluted to a 0.2-0.5% parasitemia at a 4% HCT.

### 2.4 Growth assay

Unless otherwise indicated, three replicates for every condition were grown in parallel to provide technical replicates; the replicates were grown in the same blood. Every two cycles, the samples were split into new blood (at least 50%), ensuring blood from three donors was tested over the course of the experiments, providing further replicates. Every day at ^∼^3 hours after the initial invasion window the parasitaemia was measured using flow cytometry. 5 *µ*l of each well was diluted in 45ul of PBS in a 96-well plate. The samples were stained using SYBR Green I (Invitrogen, Paisley, UK) at a 1:5000 dilution for 45 mins at 37 ° C. The samples were then run on an Attune NxT acoustic focusing cytometer (Invitrogen) with SYBR green excited with a 488 nm laser (BL1-A) and detected by a 530/30 filter. The parasitemia was measured using the Attune NXT software, for each data set, a plot of SSC-A vs FSCA was used to gate for roughly the size of an RBC, Fig. S3a, and then singlets were gated for in an FSC-H vs FSC-A plot, Fig. S3b. Next, histograms for the BL1-A intensity were used to gate for the SYBR green positive RBCs, the percentage of positive RBCs represents the percentage of infected RBCs, Fig. S3c. Every 48 h, when the culture was ring stage, it was split to give 0.5% parasitemia.

### 2.5 Shaking conditions

Shaking was performed at 45 rpm, 90 rpm or 180 rpm using an orbital shaker (Celltron from Infors HT), which has a 25 mm shaking throw. Comparison of different culture vessels was performed at 4% HCT using a: 50 ml Polystyrene tissue culture treated flask with an area of 25 cm2 (ref 353014 Falcon) with a 5 ml culture volume, a 6-well plate with a 5 ml culture volume, a 24-well plate with a 1ml culture volume or a 96-well plate with a 100 *µ*l culture volume. Different HCTs were compared in a 6-well plate with a 5 ml culture volume.

### 2.6 Calculating growth

Growth was measured by calculating the Parasite Erythrocyte Multiplication Rate (PEMR). PEMR was defined by [31] as “The ratio between the number of parasites at any point in the Intraerythrocytic development cycle (IDCn) and the number of rings in the IDCn+1.”. Custom Python scripts were used for further analysis (available at 10.5281/zenodo.13372478). As shown previously [14, 32], starting parasitemia can affect growth rate demonstrated by the negative correlation seen when the parasitemia in the initial intra-erythrocytic developmental cycle (IDCn) was plotted against the PEMR of the subsequent cycle (IDCn+1) were plotted, Fig. S4. While the target starting parasitemia for all cycles was 0.5%, splitting inaccuracies meant that there was some variation (64.9% of measurements across all assays had a parasitemia 0.25%-0.75%). The effect is most noticeable when the starting parasitemia is above 1%, therefore, any wells with IDCn¿1% were removed from the data sets (9.7% of measurements had an IDCn¿1% across all assays). Subsequent cycles were unaffected if parasitemia was correctly split to IDNn ≺1%. Plots of the PEMR by cycle were assessed by repeat and any data set removed where a very high parasitemia (typically IDCn¿5%) caused subsequent repeats to be very low (2.1% of measurements had an IDCn¿4% across all assays). These wells were manually removed by looking at the PEMR over repeats for every culture tested. The multiple invasion rates were assessed in the ring stage cultures by first using FlowJo (Tree Star, Ashland, Oregon) to plot BL1-A intensity and measured the proportion of reads in peak, Fig. S3.

### 2.7 Data availability

All code, raw data, graphs, and videos supporting the findings of this study are available in the Zenodo repository at 10.5281/zenodo.13372478.

## 3 Results

### 3.1 Significant agitation is required to prevent RBCs from sedimenting

We began by exploring fluid motion in standard malaria parasite culture conditions - a 6-well plate with each well containing a 5 ml culture volume at 4% haematocrit (HCT) (i.e. 4% w/v RBCs/medium). To understand the conditions that the cells are exposed to in these wells, we filmed how the fluid motion of blood changed as the shaking speed increased, Movie S1 and Fig. S1. At static and low rotational speeds, the RBCs remain sedimented at the base of the well. As rotation speed increases the media begins to move above the sedimented RBCs, which results in the cells sedimented at the base of the well being subject to the wall shear stress (WSS). Then, as a critical threshold is surpassed, which we term the *critical clustering transition*, blood begins to cluster in the centre of the well, forming a toroidal vortex of RBCs. Further speed increases cause a corresponding increase in the motion of the vortex structure, and more RBCs become suspended in the laminar flow region surrounding the central vortex. As orbital rotation speed increases further, the RBCs gradually get better mixed into the bulk media, and the flow transitions to become more turbulent; eventually, all the RBCs are in suspension. In this culture format, three shaking speeds can capture the different fluid dynamic conditions: 45 rpm, where the fluid is moving over the sedimented blood but the RBCs are not moving, 90 rpm which is just above the critical clustering speed and the RBCs have largely moved into a central vortex but have not yet dispersed into the surrounding medium, and 180 rpm where the conditions are highly turbulent and the RBCs are completely mixed through the medium, Fig. 1a.

**Figure 1.**
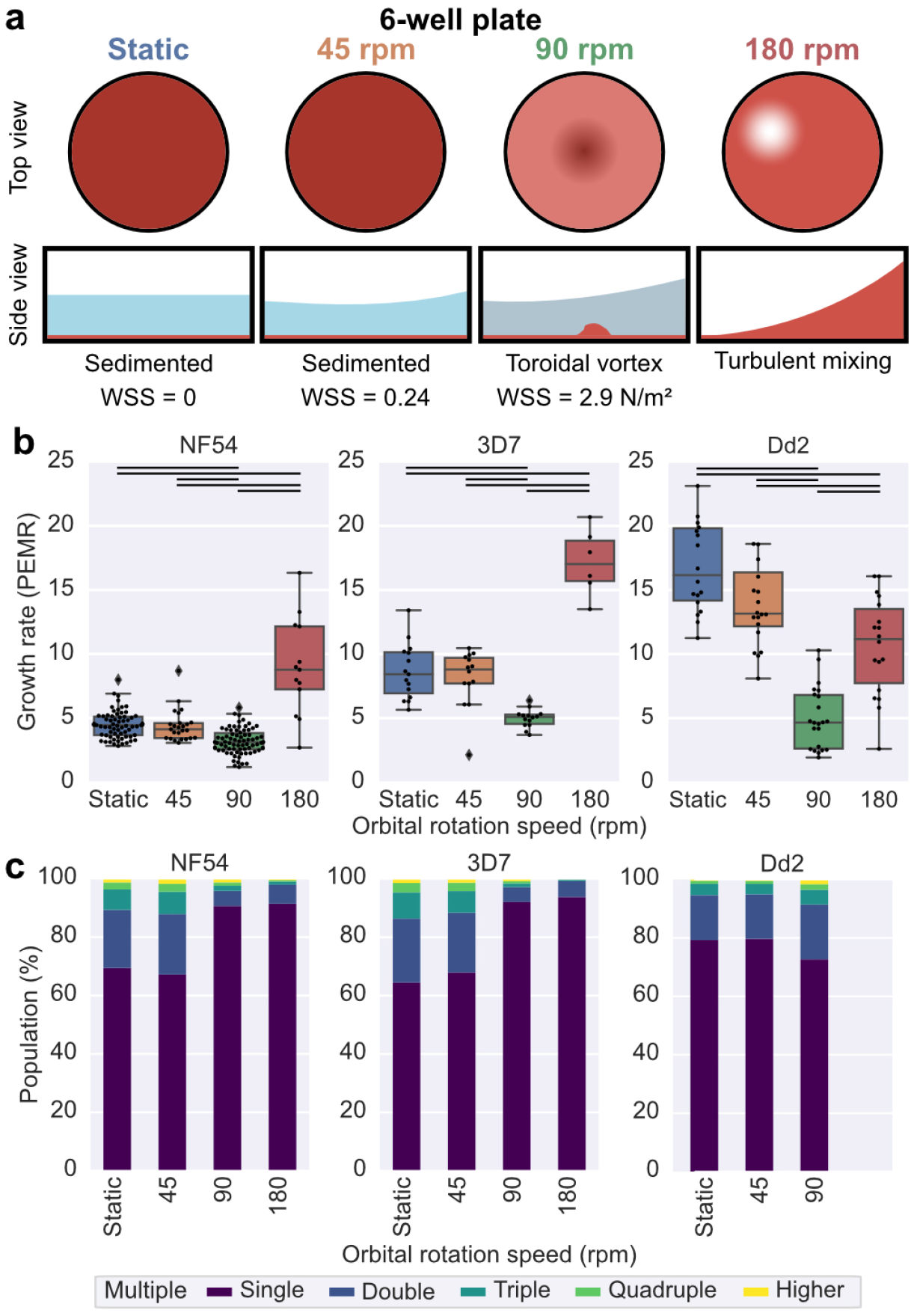
Shaking speed affects growth rate of wild-type *P. falciparum* lines: NF54, 3D7 and Dd2. All data was collected for samples cultured in a 6-well plate, 5ml culture volume and a 4% HCT. **(a)** A schematic to represent visually what the motion of the blood looks like at the conditions for which growth was compared. At static, the blood is sedimented at the base of the well; at 45 rpm the fluid just starts to move over the sedimented blood; 90 rpm is just above the critical clustering speed and the RBCs are largely confined to the vortex, and at 180 rpm the conditions are highly turbulent and the RBCs are completely mixed through the medium. The simulated mean WSS are shown for the conditions where the blood is still sedimented. **(b)** The growth rate is shown as the Parasite Erythrocyte Multiplication Rate (PEMR). The data displayed is from all experiments run under the indicated conditions across multiple assays. The lines between conditions indicate the significantly different conditions (t-test) at a greater than 5% significance level. **(c)** Graphs represent the mean frequency of different numbers of parasites in a single RBC for a given speed for each wild-type line. The frequency of single-infected, double-infected, triple-infected, quadruple-infected, and higher-infected rings is shown as a fraction of the total infected ring-stage parasites. The number of rings per erythrocyte was measured using the peak intensity of the ring-stage infected samples and the number of counts in each peak. Data sets without clearly defined peaks were excluded, which is why there is no data for Dd2 at 180 rpm.

Quantifying the WSS in these round wells under orbital motion is difficult because the fluid motion is non-uniform [17]. To estimate WSS, we used the model presented in [8] and based on [33, 34]. We simulated the mean WSS over time at three points along the bottom of the well (25%, 50%, and 75% between the wall and centre) and reported the mean of these, Fig. 1a and Fig. S2d. WSS increases with orbital rotation speed. For the 6-well plate, the mean WSS at the clustering transition speed (85 rpm) was simulated to be 2.7 N/m^2^, which interestingly is comparable to measurements of WSS in human microvessels [35], were late stage parasites sequester [4].

### 3.2 Shaking can both decrease and increase the growth rate of wild-type *P. falciparum* parasites

The growth of three wild-type *P. falciparum* strains (NF54, 3D7 and Dd2) were compared at these three different shaking speeds while keeping all other conditions constant (6-well plate, 5 ml culture volume, 4% HCT). Growth was defined using the Parasite Erythrocyte Multiplication Rate (PEMR), as defined by [31], calculated by dividing parasitemia after invasion (ring stage culture) by parasitemia the day before (trophozoite/early-schizont stage culture). Parasitemia was measured using flow cytometry to calculate the percentage of DNA-positive (i.e. infected) RBCs, Fig. S3. The effect of starting parasitemia was carefully controlled for, Fig. S4. The wild-type lines have different PEMRs under static conditions. NF54, which originates from a patient in the Netherlands although genomic analysis suggests it is likely of West African origin [36], had a growth rate PEMR of 4.2 ± 0.2. 3D7, Fig. 1a. 3D7 is a clone of NF54 [36] that has been cultured independently for 30 years and while being genetically very similar [37] has multiple diverged phenotypes [38, 39, 40] including shown here in growth rate with a PEMR of ± 0.6, Fig. 1b. Dd2 is a strain of Southeast Asia origin that was chosen for comparison as it has many invasion characteristics distinct from 3D7/NF54 and it had the highest growth rate with a PEMR of 13.7 ± 0.5, Fig. 1b. These measurements are comparable with previous measurements of growth with qPCR in which 3D7 had a mean growth rate of 8.1 and Dd2 10.5 (assays performed at 3% HCT and 0.02% starting parasitemia) [41]. For all three wild-type lines, there was no significant difference in PEMR between static and 45 rpm, where the RBCs are not in suspension. At 90 rpm, just above the critical clustering speed, the growth rate drops for all lines, while at 180 rpm, the growth rate increases for 3D7 and NF54 and decreases relative to static for Dd2. This shows that shaking does not always produce better growth/invasion rates and that the relationship between growth and shaking is complex.

We quantified the degree to which erythrocytes were infected with multiple parasites based on peaks in the intensity of DNA staining in the ring stage cultures as described previously by [42], Fig. S1d. This was possible as all assays were set up using parasites tightly synchronised to a 3 hour window and measurements were taken at the ring stage, before parasites began DNA replication (see Methods for details). As reported previously, we refer to the number of rings measured in each RBC as its multiplicity of infection (MOI), [43]. Previous observations using Giemsa smears have shown up to five rings can be observed in a single RBCs and that different strains have different rates of mean MOI rates [43]. For 3D7 and NF54, there was no difference in the proportion of RBCS with two or more rings between static and 45 rpm, but much lower rates at both 90 rpm and 180 rpm, Fig. 1c, in keeping with previous reports of lower rates of mutiple infected rings when culturing under shaking conditions [15, 16, 11, 12]. This trend was not observed for Dd2, Fig. 1c, but the MOI data for Dd2 is less reliable as its replication cycle is shorter. It is important to note that MOI and growth may be linked, but are not definitively so - despite 90 rpm and 180 rpm having opposing effects on growth for 3D7 and NF54, both speeds reduced mean MOI rates.

### 3.3 The critical clustering speed correlates to a drop in growth rate across different culture conditions

To further explore the complex relationship between growth rate and shaking conditions, we utilised the fact that changing vessel shape and size changes the resultant fluid flow produced at a given speed. We performed the same growth assay in three additional vessels, Fig. 2a. Filming the changes in the blood motion across shaking speeds for these new conditions showed a consistent fluid flow change as the rotational speed was increased, Movie S2-S4 and Fig. S5. However, the smaller the well, the higher the orbital rotation speed required to reach the critical clustering transition. This pattern was observed for all vessels, but the fluid flow was more irregular in the culture flask due to its non-uniform shape, leading to more fluid mixing even at very low shaking speeds. We tested the growth in the different vessels of wild-type lines NF54 and Dd2 at the same range of shaking speeds tested previously, Fig. 2a. The critical clustering transition speed (when blood began to cluster in the centre of the well, see Fig. S2a) provided a way to normalise the shaking speeds across round well sizes. When the mean growth rates were plotted relative to this normalised orbital rotation speed, there was a clear pattern for both wild-type NF54 and Dd2, with the dip in growth rate occurring close to the clustering transition speed, and growth increasing once normalized rotation increases beyond this transition, Fig. 2c and S2c. Interestingly, simulating WSS for the different well types shows that the shear forces at the critical clustering transition speed are similar in magnitude, Fig. S2d. This is likely because the shear stress is a determining factor in suspending the RBCs. Thus, normalizing to the critical clustering speed will, to some degree, normalize the shear forces experienced by the parasites.

**Figure 2.**
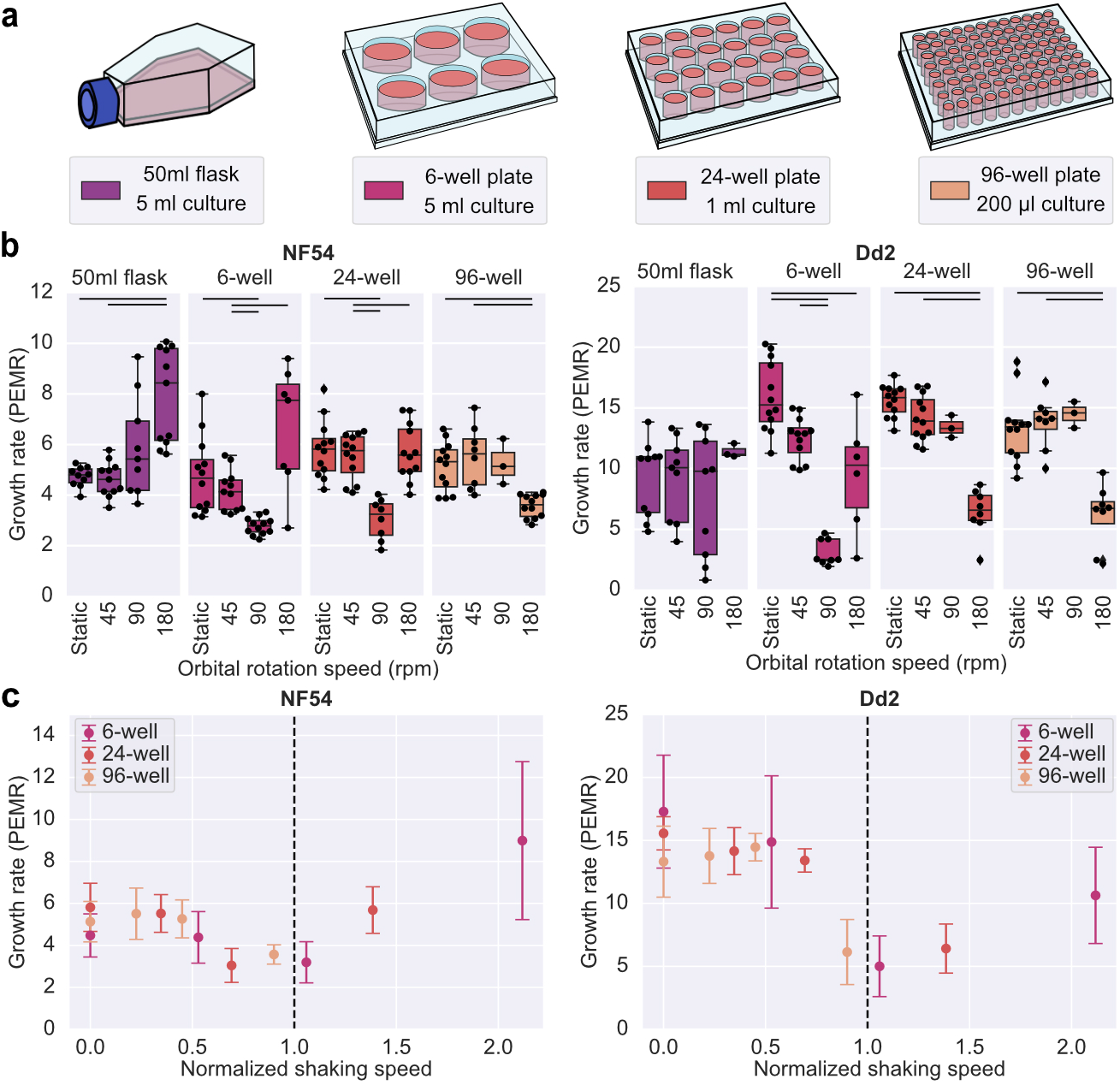
The type of culture container affects the growth rate of wild-type lines of *P. falciparum*. **(a)**Schematics illustrate the culture conditions that were compared. **(b)** The wild-type lines NF54, 3D7 and Dd2 were tested. All cultures were kept at a 4% HCT. The mean growth rate is shown as the parasite erythrocyte multiplication rate (PEMR). The experiment was performed in triplicate technical repeats; each point shows a PEMR measured for a single invasion cycle for an individual well. The lines between conditions indicate the significantly different conditions (t-test or rank sum) at a greater than 5% level of significance. **(c)** Growth rates from (b) for the round wells plotted against the normalised shaking speed. The shaking speed was normalised to the rotation speed at which the blood was first seen to cluster in the centre of the well, Fig. S2a.

As expected, the culture flask/volume also changed how a given shaking speed affects the multiple invasion rates, Fig. S6a. The switch from high to low MOI rates also corresponds to the clustering transition speed for NF54, Fig. S2b. At this point, the blood starts to move into suspension, reducing the probability that two merozoites that egress from a given schizont will contact and invade the same RBC. Another variable likely affecting how shaking impacts growth is the erythrocyte density in the culture (haematocrit). The growth of 3D7 and NF54 was compared at 2%, 4% and 8% HCT, Fig. S2a. Altering haematocrit changes how a given shaking speed affects growth but not the multiple invasion rate, Fig. S2b. However, in general, a comparison of a single haematocrit as shaking speed increases is consistent with the pattern of decreased growth followed by increased growth. This demonstrates that it is important to keep haematocrit consistent when making comparisons between growth rates under shaking conditions.

### 3.4 Deletion of PfEBA and PfRH invasion ligands changes the impact of shaking on growth

We hypothesised that high WSS leads to lower growth rates because it makes it harder for merozoites to attach to RBCs during invasion. Many members of the PfEBA and PfRH family of ligands have linked to the initial steps of invasion [44], and we recently showed several members of these families are key determinants of the strength of attachment between merozoites and RBCs, [45]. We were therefore interested in whether disruption of PfRH and PfEBA proteins would impact the change in growth between static and shaking conditions at the critical transition point (6-well plate at 4% HCT and 90 rpm). A panel of PfRH and PfEBA knock-out lines in the NF54 background were tested along with two control knock-out lines constructed in the same manner [45]. The control lines have the gametocyte-specific proteins PfP230P or Pfs25 knocked out, both proteins that have no known role in invasion [46, 47]. Wherever possible, two clones (cX, cY) were tested of a given knock-out to assess the variability between clones. The data for the static growth rates of these lines was previously published [45]. As previously reported by both us and others [26, 48, 27, 49, 50, 51], deletion of these ligands, which are redundant and largely thought to be functionally interchangeable, had no impact on growth under static conditions, although there were some minor differences in growth relative to wildtype NF54 in some knockouts. These minor differences are likely due to epigenetic differences acquired in growth rates or invasion phenotypes of clones during the extended period of culture required to generate transgenic parasites, as has been reported previously [52, 53, 32]. Several of the lines showed higher rates of multiply infected erythrocytes than NF54, including ΔPfs25c1/3, ΔPfEBA181c1/2, ΔPfRH2ac1 (but not c3) and ΔPfRH4c1, Fig. S7.

When PEMR was compared between growth in static conditions and shaking at the critical clustering transition point for the wild-type NF54 line and control lines (ΔPfP230P and ΔPfs25), there was little (less than 30%) or no significant difference, Fig. 3, as expected. By contrast, several PfEBA and PfRH knockout lines had a significantly lower PEMR when grown in shaking conditions at the critical clustering transition point compared to static conditions: ΔPfEBA140c3/c4 (c3 2.4-fold lower p = ≺0.0000, c4 2.6-fold lower p = ≺0.0000), ΔPfEBA175c6 (c2 3-fold lower p = 0.0003, c3 2.5-fold lower p = 0.0013), ΔPfRH1c1 (1.7-fold lower p = 0.0009) and ΔPfRH4c1 (1.7-fold lower p = ≺0.0000), Fig. 3. It is interesting that for the ΔPfEBA181c1/c2 while there was no significant difference in PEMR between static conditions and 90 rpm; at 90 rpm, the PEMR for ΔPfEBA181c1/c2 was significantly higher than NF54. This suggests that growth under shaking at the critical transition point can reveal growth/invasion phenotypes that are not detectable through static growth assays. In addition, the loss of several of the PfEBA and PfRH proteins (PfEBA140, PfEBA175, PfRH1 and PfRH4) seems to be more detrimental to growth under these shaking conditions, meaning these proteins may be more important to binding in conditions of high WSS, which in this assay set up are similar to the conditions that schizonts will experience in the microvasculature.

**Figure 3.**
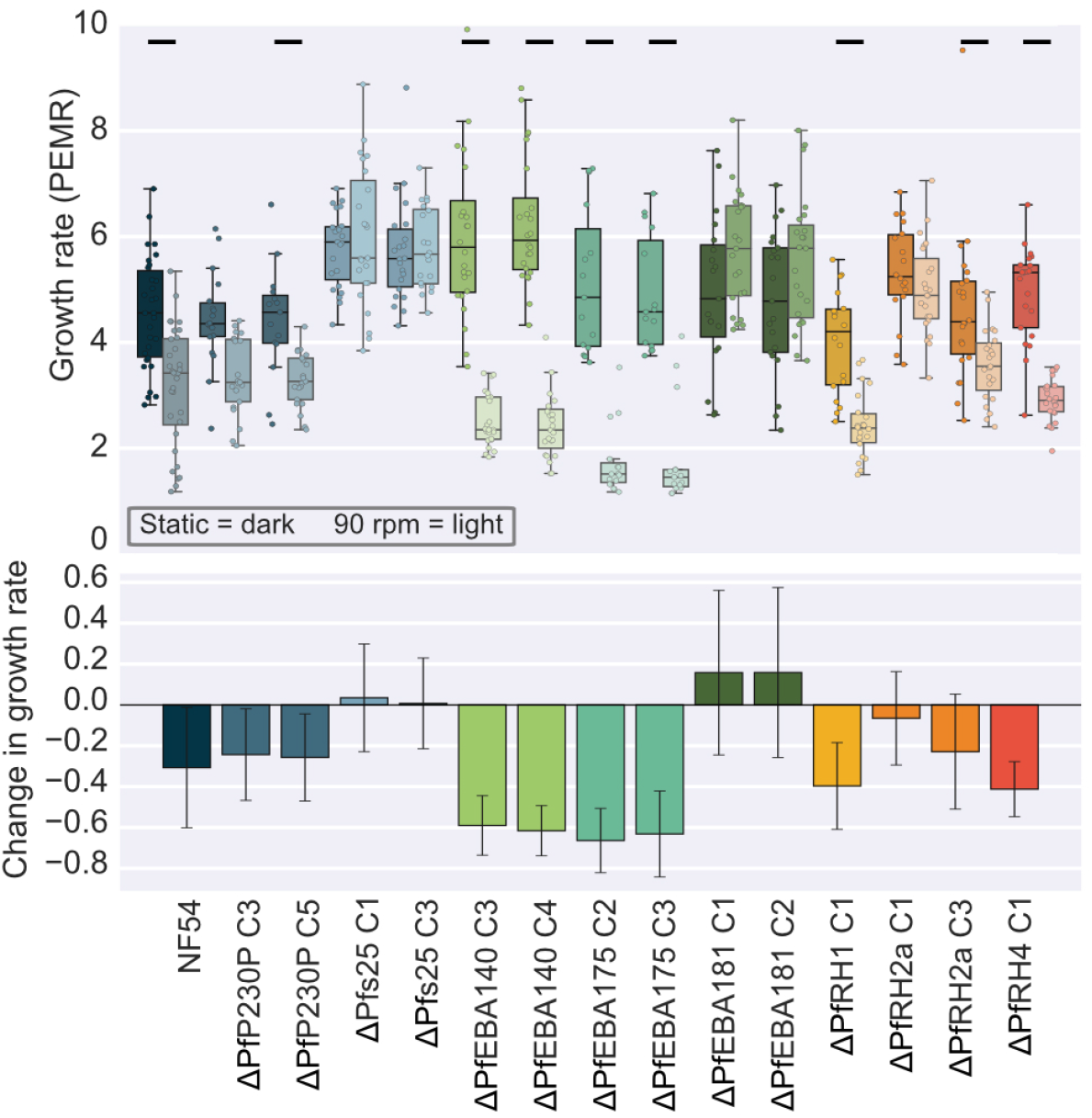
Knocking out specific invasion ligands shows flow-dependent effects. Experiments are carried out to compare PfEBA and PfRH knock-out line growth rates under static and detrimental shaking conditions, measuring in static and at 90 rpm; two clones of each line were tested wherever possible. All cultures were grown at 4% HCT in 5 ml in a 6-well plate. The top box plot shows the mean Parasite Erythrocyte multiplication rate (PEMR) of the knock-out lines. The central red line shows the median, with the top and bottom of the box at the 25th and 75th percentiles and the whiskers showing the total range of the data. The bottom bar plot shows the relative difference in growth in shaking compared to static for each line. The error bars show the standard deviation.

### 3.5 Changes in growth rate are predominantly due to changes in invasion

Finally, we tested whether the changes seen in growth were due to the effects of shaking on invasion itself, rather than effects on some other point in the 48-hour life cycle. Schizonts were isolated from a tightly synchronised culture, resuspended at 4% HCT and then split across the wells of 6-well plates. The plates were then subjected to either static or shaking conditions for 3.5 hours to allow for invasion to occur. The percentage of rings (post-invasion) and schizonts (pre-invasion) was measured for each well before and after this 3.5 hour window using flow cytometry; the invasion rate was then calculated by dividing the change in the newly invaded RBCs (rings) by the change in the late-stage infected RBCs (schizonts), Fig S9a. Invasion rate showed the same pattern as growth rate, decreasing at 90 rpm and increasing at 180 rpm. To allow comparison between conditions, results were normalised to static, Fig S9 b. There was no significant difference in the change in schizonts between conditions, although the low percentage of schizonts in the sample meant measurement accuracy was low, causing large variations in measurements. Fig S9 c shows no correlation between change in schizonts and corresponding growth rate (Pearson correlation coefficient -0.09), meaning the changes in invasion rate are not due to shaking affecting the ability of parasites to egress from the schizonts. However, the change in rings and invasion rate correlate highly with the previously measured change in growth (Pearson correlation coefficient 0.94 and 0.84, respectively), Fig S9 c, for all lines and conditions tested. This indicates that the effect of shaking on growth is predominantly due to its impact on invasion, as predicted by the effect of deleting individual invasion ligands on growth under shaking conditions.

## 4 Discussion

By growing *Plasmodium falciparum* parasites on rotational platforms shaking at different speeds and in different culture flasks and volumes, we show for the first time an interesting behaviour of decreased parasite growth at low orbital rotation speeds and then increased growth under higher speeds. While at first glance perhaps unexpected, calculation of the forces that the parasites are exposed to show that these changes can be explained by known elements of fluid mechanics. Fluid motion generated by the orbital shaker creates a non-uniform shear stress field even in circular wells [17] and observation of the changes of RBC behaviour in fluid motion showed that different phases can be distinguished as shaking speeds increase, as described above and shown in Fig. S1. The speed at which the RBCs transition from sedimented to beginning to cluster in the centre of the well, which differs depending on vessel size/geometry and culture volume, represents a point at which the fluid motion will exert high sheer stress on the sedimented blood. Strikingly, regardless of strain or vessel used, this point also resulted in the lowest growth rates. The fact that changes in growth rate at different shaking speeds are mirrored almost excatly by changes in invasion rate at those speeds, Fig S9, suggests strongly that this dip in growth occurs because the high WSS at this transition point reduces the rate of successful attachments between merozoites and RBCs. We expand on that finding by showing that disruption of several *P. falciparum* ligands known to be involved in attachment (PfEBAs and PfRHs), further reduce growth rates at this critical transition speed, Fig. 3. At higher shaking speeds, when the blood starts to become suspended and as the conditions become more turbulent, growth rates increase and the MOI rate falls, Fig. 2 and S6. These turbulent conditions likely increase the frequency of contact between merozoites and RBCs, which overcomes the detrimental effect of reduced successful attachments due to the increased WSS. We have previously shown that different wild-type lines have different attachment frequencies and detachment forces using an optical tweezer-based assay [45], which would be one reason why different lines are affected to different degrees by a given shaking condition, as shown in Fig. 1 and in existing literature [9, 13, 54]. This relationship is summarised in Fig. 4.

**Figure 4.**
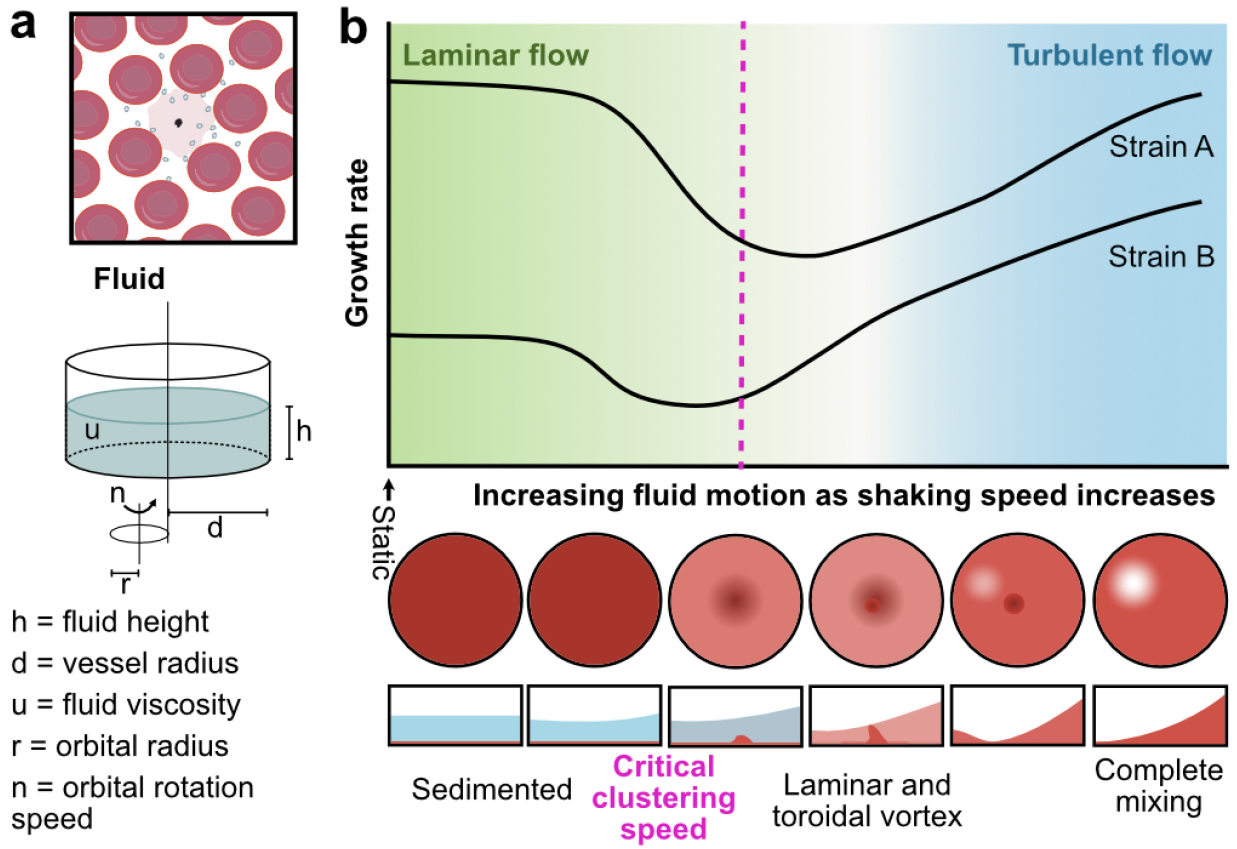
Summary of effects of shaking on *P. falciparum* growth. **(a)** Summarize the factors that determine fluid motion when a round well is on an orbital shaker. **(b)** Shows a cartoon of the relation we have shown between increased orbital rotation speed and growth in round wells. Alongside are the observed changes in a fluid motion.

Our systematic testing of multiple conditions shows that multiple factors must be considered when culturing *P. falciparum* under shaking conditions, particularly the geometry of the culture vessel, Fig. 4a. *P. falciparum* grows very differently while shaking at different orbital speeds, Fig. 1, in different flasks, Fig. 2a, and over different haematocrits, Fig. S8. For example, no drop in growth in shaking relative to static conditions was observed when culturing in the square flasks under the conditions shown in, Fig. 2a. This is likely because square flasks produce more mixing than the round wells even at low speeds, Fig. S1, meaning either the lowest speed tested (45 rpm) was too high to create a toroidal vortes or the shear forces at the critical clustering speed may be too low to impact the growth rate. This data is in keeping with previous studies where culture conditions are detailed. Where increased growth was reported, flasks were used to grow cultures [16, 11, 12, 13]). Similarly, the previously published example of decreased growth in shaking conditions was performed using a 2 ml culture in a 12-well plate at 125 rpm [14], which based on correlation with our normalised rotational speed data, was likely close to the critical clustering point, Fig. 2c and S2a. The data and normalisation presented here therefore provide a clear framework to design future experiments which can be adapted to the specific set-up in each lab, as it is easy to visually determine the lowest orbital rotation speed at which blood starts to cluster in whatever round, flat-bottom well plate is being used. This provides an easy way of determining conditions that are likely to produce either detrimental (at the critical clustering point) or favourable (above this transition point) growth conditions.

The method presented here also provides a new and simple phenotyping assay to assess factors important to growth and invasion under physiologically relevant shear force conditions. As described above, for a 5 ml culture in 6 well plates or 12.5 mm radius filled with 4% HCT culture and shaken at 90 rpm, the mean shear was 2.5 N/m^2^, while WSS will vary across the vessels of the circulation system, this is similar to the mean WSS measured in human conjunctival capillaries at 1.5 N/m^2^ [35]. As schizonts have been shown to sequester in the microvasculature [4], this means that the forces experienced by parasites at the critical shaking speed is in the physiological range.

Using this new phenotyping assay in combination with PfEBA and PfRH knockout lines, we have established that the invasion ligands PfEBA140, PfEBA175, PfRH1 and PfRH4 are more important for growth under conditions of high shear stress than in static conditions, where none of these lines had growth defects relative to controls [45]. In the case of PfRH4, this fits with the previous observation that PfRH4 expression is up-regulated in some lines when they are grown under constant shaking conditions [13, 54]. However, this specific phenotype, of decreased invasion in the absence of PfRH4, has not previously been observed, and nor have the other ligands been linked to invasion under shaking conditions. When the attachment characteristics of the same panel of PfEBA and PfRH knockout lines were assessed using an optical tweezer-based assay, the only line in which there was a significantly decreased detachment force was ΔPfRH4 [45]. This all points towards the strong attachment of PfRH4 being important to efficient invasion under shaking conditions. It is currently thought that the interactions of PfRHs and PfEBAs during invasion are largely redundant, and this data provides evidence that several proteins may actually be more crucial under high shear forces similar to those experienced *in vivo*. An area for future investigation is whether different ligands could be utilised in different environments in the body, which motivates the further development of a system where invasion can be followed under a more uniform flow that can be more easily quantified and controlled.

## Acknowledgements

This research was supported by the Cambridge Institute of Medical Research (CIMR) Flow Cytometry Core Facility, and we wish to thank Reiner Schulte, in particular, for advice and support in flow cytometry. We would also like to thank Theresa Feltwell, Nadia Cross and the CIMR and Physics of Medicine Department support staff for their invaluable logistical support. This work was funded by Wellcome, grant numbers (220266/Z/20/Z to JR) and (222323/Z/21/Z to EK); EU-EC Marie Curie ITN PyMot (955910 to MK) and Engineering and Physical Sciences Research Council (EPSRC) grant (EP/W004453/1 to PC). For the purpose of open access, the author has applied a CC BY copyright license to any Author Accepted Manuscript Version arising from this submission. The funders played no role in the study design, data collection and analysis, decision to publish, or preparation of the manuscript.

