## Supplementary figures for "Under conditions of high wall shear stress, several PfEBA and PfRH ligands are important for *Plasmodium falciparum* malaria blood-stage growth"

### Supporting Information for

#### Growing Malaria Parasites at a Critical Shaking Speed Mimicking Physiological Flow Reveals New Phenotypes for Invasion Ligands

**Emma Kals, Morten Kals, Pietro Cicuta, Julian C. Rayner**

**Julian C. Rayner.**
****

##### This PDF file includes:

Figs. S1 to S9
Legends for Movies S1 to S4
SI References

##### Other supporting materials for this manuscript include the following:

Movies S1 to S4

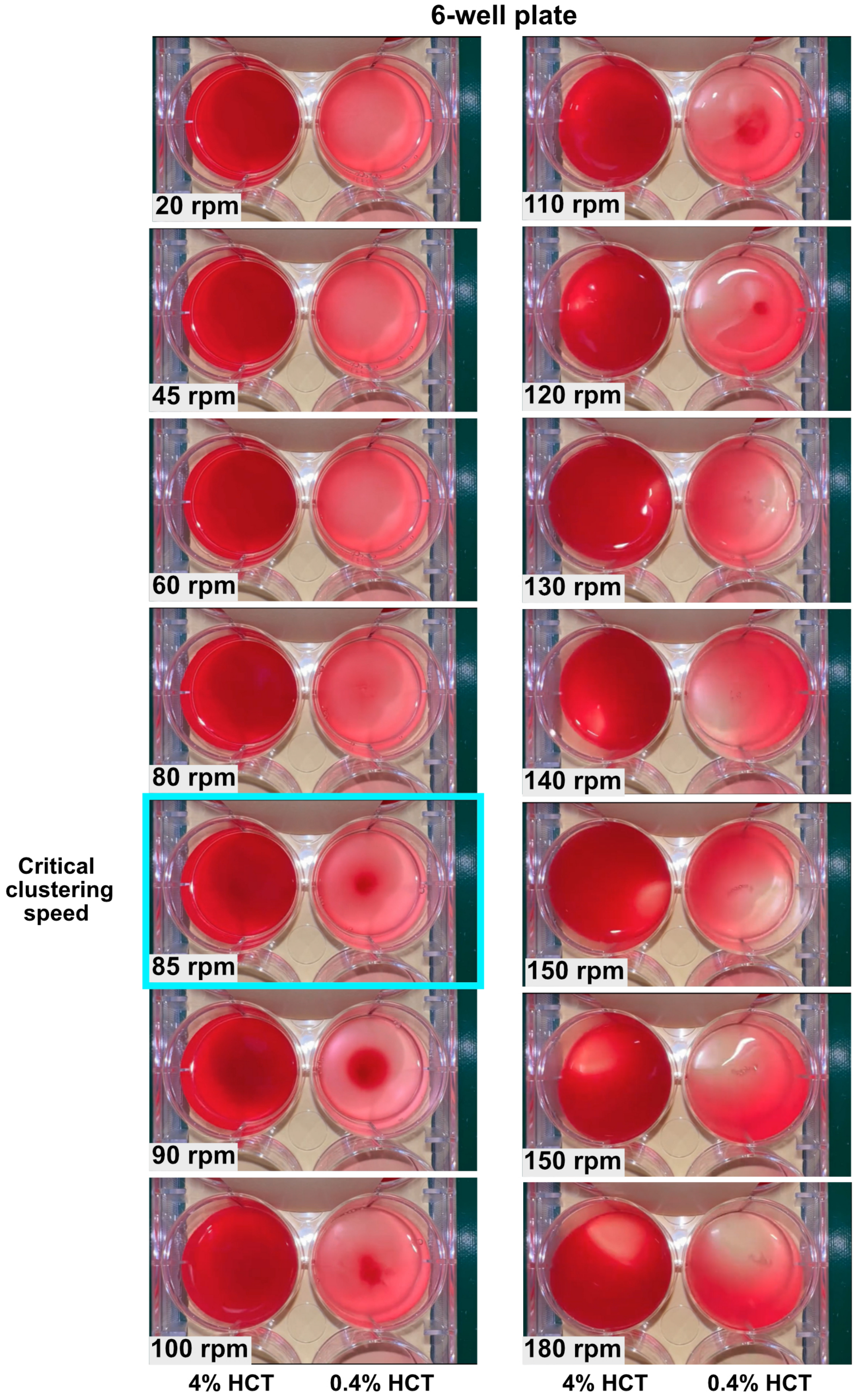


**Fig. S1. Comparison of the effect of different shaking speeds on the suspension of blood in culture.** Data is shown for 6-well flat-bottomed well plates with 5 ml culture volume at 4% (left well) and 0.4% HTC (right well). Images show how the blood goes from sedimented uniformly at low speeds, to clustering in the center, to being uniformly suspended in the culture. The blue outline represents the critical clustering speed where clear clustering starts to occur.

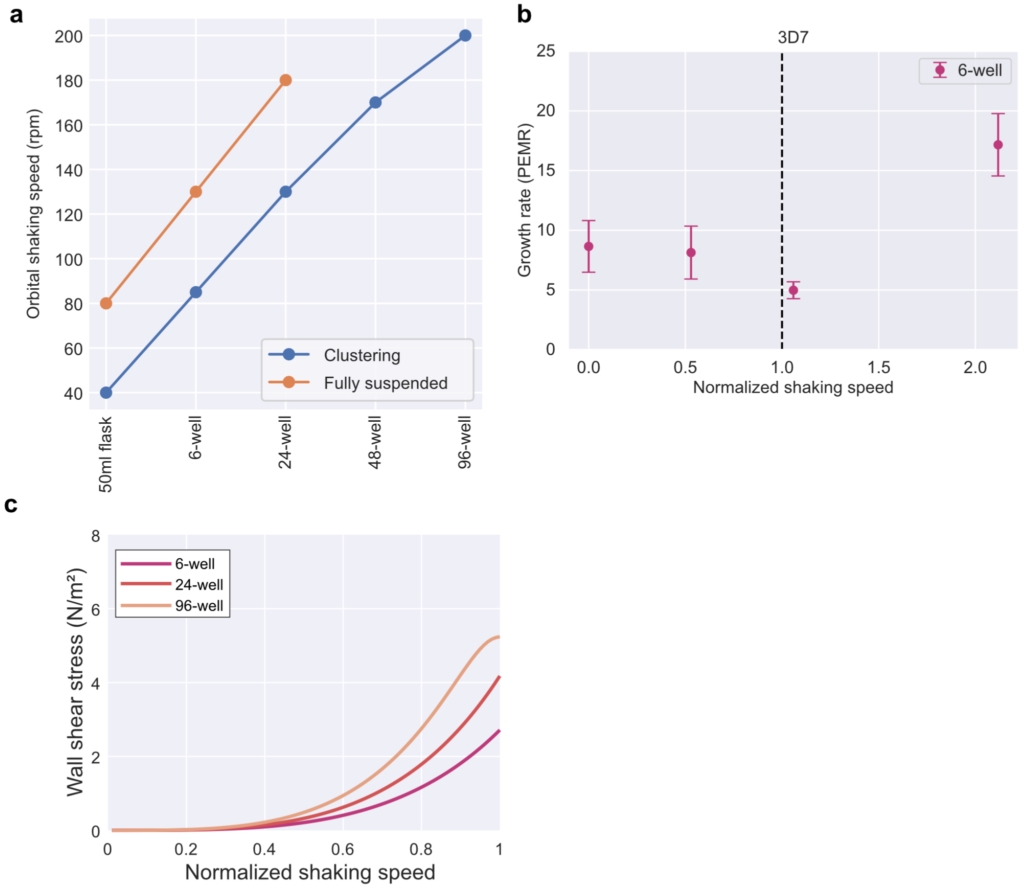


**Fig. S2. Characterizing fluid motion (a)** Shows the orbital shaking speed for our media volume at which the RBCs start to cluster in the centre of the well and at which the RBCs are fully suspended in the bulk medium. For the 48 and 96 well plates, full suspension does not occur below 200 rpm, which was the top speed of our shakers. **(b)** Attempting to correlate the clustering thresholds with Reynolds number does not give a good correspondence. Reynolds number was calculated using the formula for round wells presented in (1) **(d)** Simulating wall shear stress (WSS) for each plate size at the volumes used in our experiments using the PT-Stokes model from (2) shows that the shear forces experienced in each plate type are of the same order of magnitude at the clustering speed threshold. Shaking speed normalized to clustering speed for each plate well size.

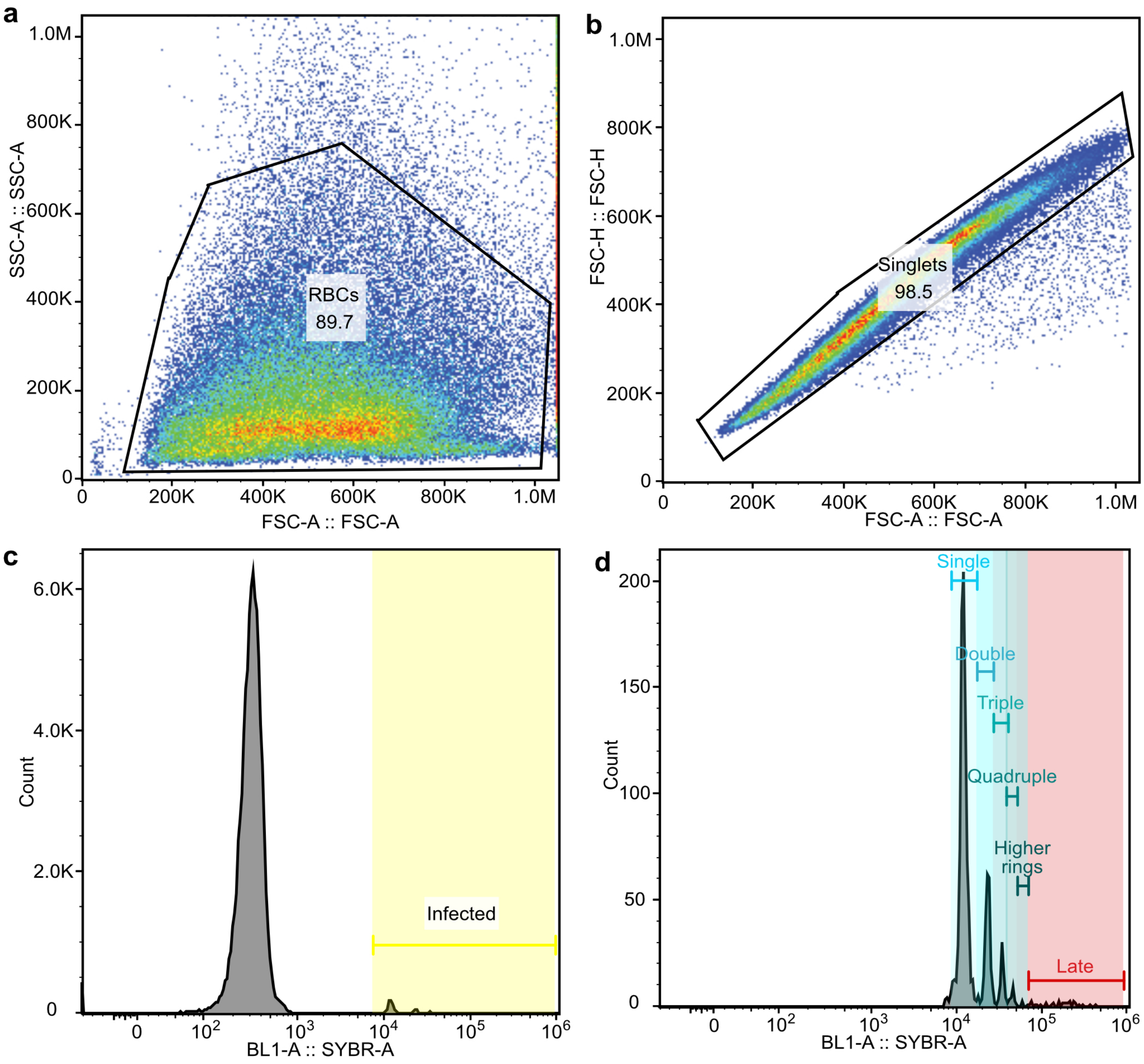


**Fig. S3. Gating used to determine parasitemia and rate of multiple infections based on flow cytometry.** Gating steps were taken to measure multiple invasion rates during the growth assay. The graphs shown are from one well on day 5 from NF54 in static condition for the growth assay that compared the PfEBA and PfRH knockouts. Analysis was done in FlowJo. The gates shown here were applied to all the data. **(a)** SCC-A vs FSC-A was plotted, and a gate was drawn to select for particles detected of roughly the right size for erythrocytes. **(b)** FSC-H vs FSC-A was plotted, and a gate was drawn to select the predominantly singlet population to exclude any signal from two or more erythrocytes clustered together. **(c)** A histogram of the BL1-A laser intensity was plotted, which detected the SYBR Green staining of the infected erythrocytes. A gate was set based on the signal of unlabelled erythrocytes. Anything with a high signal was defined as infected. Parasitemia was determined as the percentage of counts in this region out of the total number of counts after the initial two gates were applied. **(d)**Finally, for the samples collected when the parasites were early rings, additional gates were used in the region defined as infected for the peaks which represented single or multiple infected erythrocytes. This allowed calculation of the fraction of infected erythrocytes that had single, double, triple, quadruple, or higher infections and the even higher intensity region which represented any late-stage trophozoites or schizonts present.

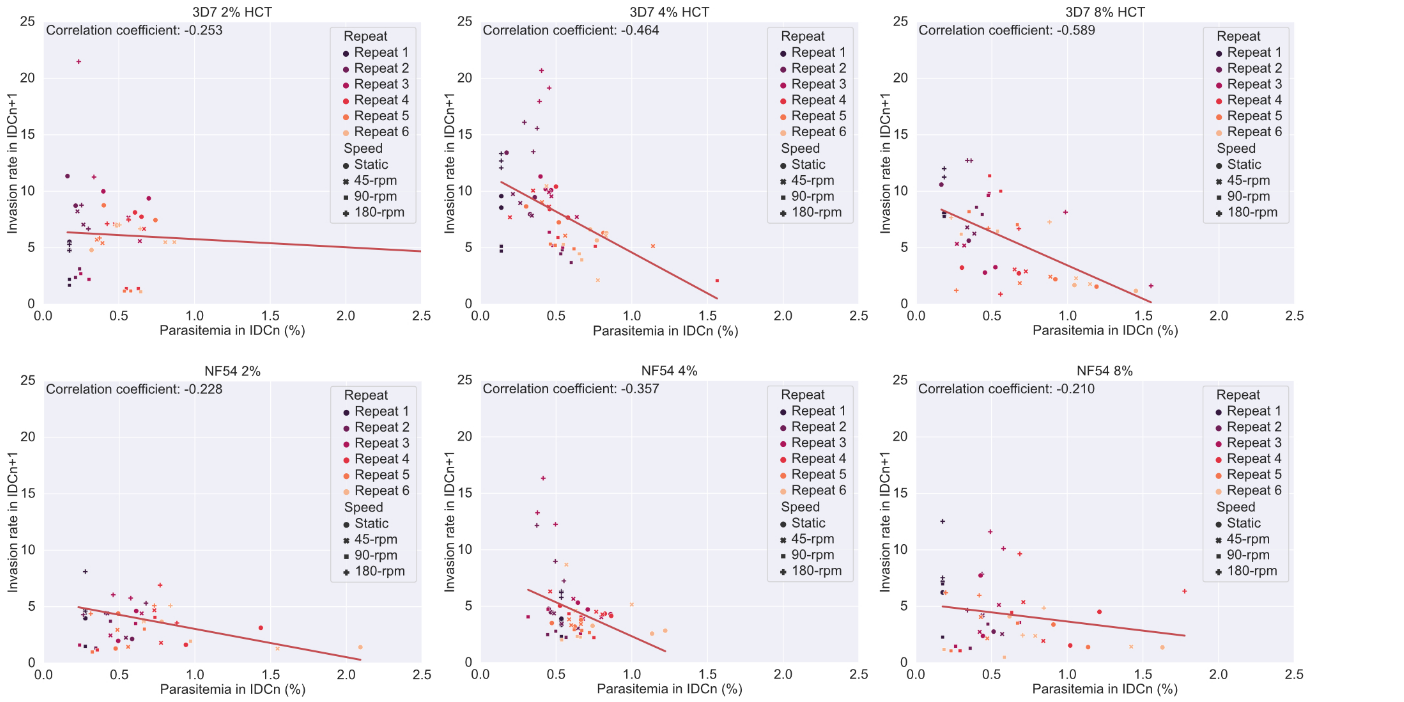


**Fig. S4. The effect of the starting parasitemia on invasion rate.** Plotted is the impact of the parasitemia in the initial intra-erythrocytic developmental cycle (IDCn) on the invasion rate of the subsequent cycle (IDCn+1). The data shown is for all NF54 and 3D7 data sets in a single assay in which the effect of different haematocrits was tested. Repeats were collected over different invasion cycles, as shown by different colours, and the different shaking speeds as shown with different symbols.

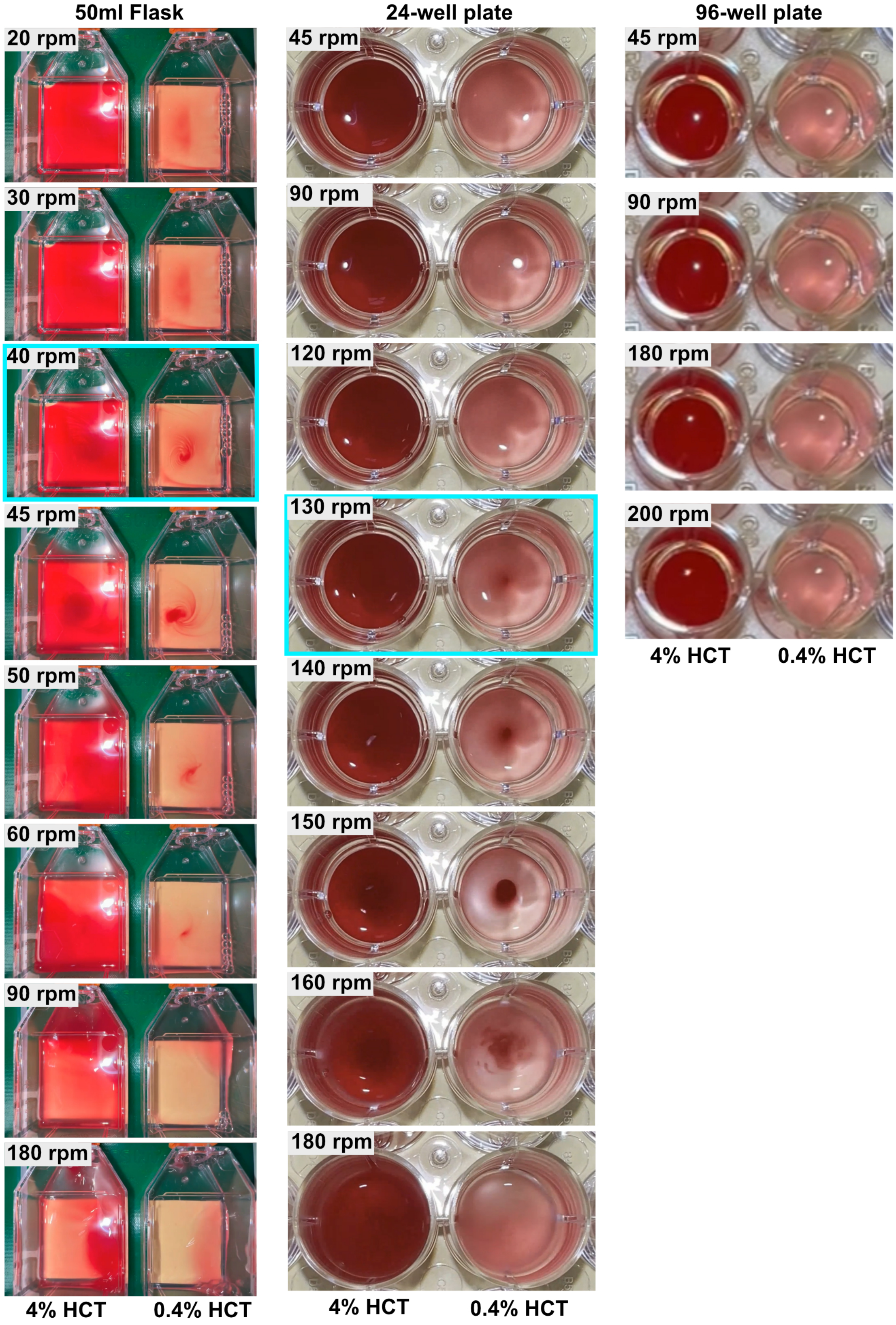


**Fig. S5. Comparison of the effect of different shaking speeds on the suspension of blood in culture in different culture vessels.** Data is shown for a 50 ml culture flask with a 5 ml culture volume, a 24-well flat-bottomed well plate with 1 ml culture volume and 96-well flat-bottomed well plate with 100 $\mu$l culture volume. All are at 4% (left well) and 0.4% HTC (right well). Images show how the blood goes from sedimented uniformly at low speeds to clustering in the center to being uniformly suspended in the culture. The blue outline represents the critical clustering speed where clear clustering starts to occur.

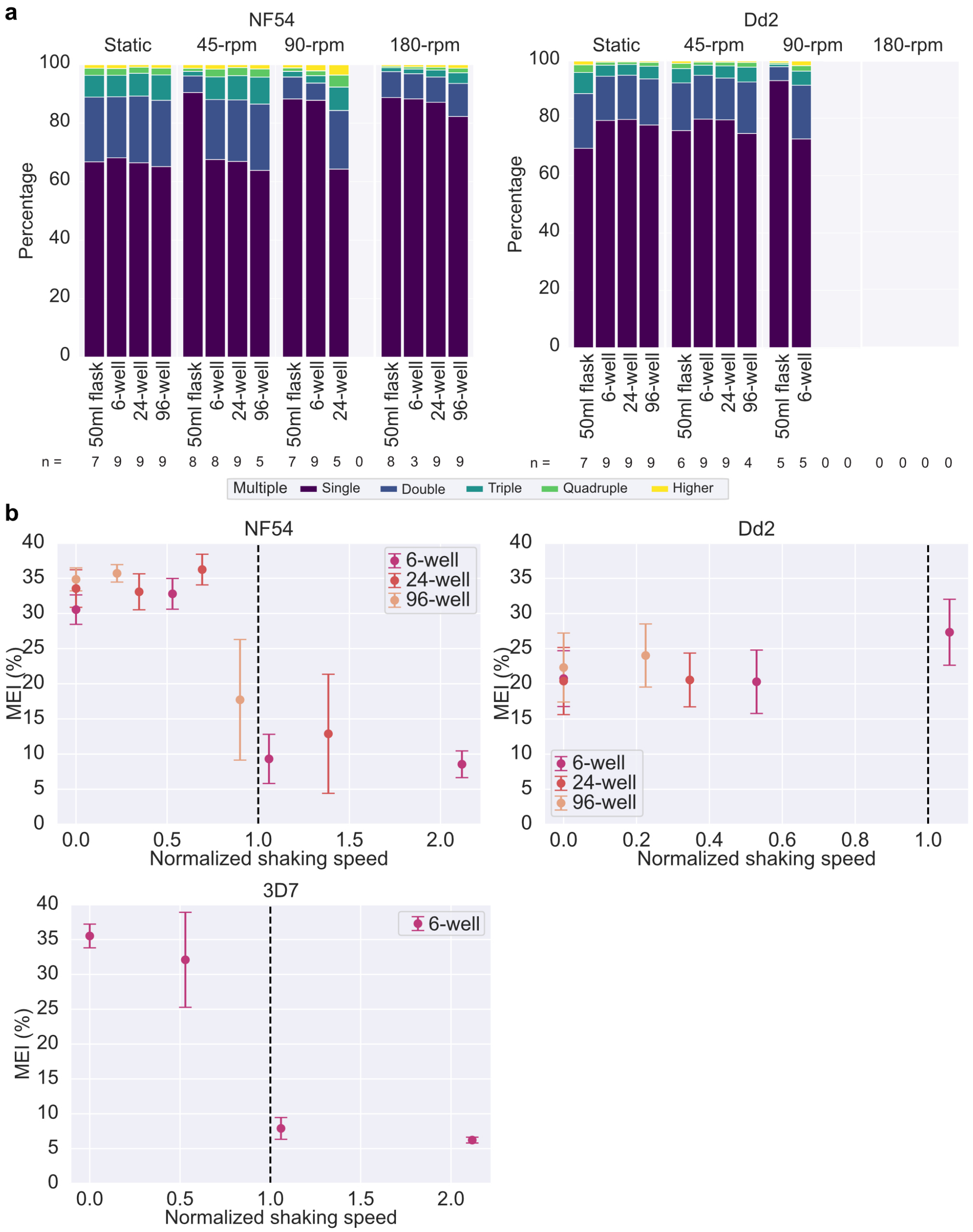


**Fig. S6. Comparison of the effect of different culture containers and culture volumes on the impact of shaking speed on multiple invasion rates.** All cultures were kept at a 4% HCT. All data presented in the figure was collected in parallel. This is the data for the same assay as in Fig 3. **(a)** Graphs representing the mean frequency of different multiple erythrocyte infections for a given speed for each wild-type line. The frequency of single-infected, double-infected, triple-infected, quadruple-infected, and higher-infected rings is shown as a fraction of the total infected ring-stage parasites. The number of rings per erythrocyte was measured using the peak intensity of the ring-stage infected samples by counting the fraction of counts in each peak. **(b)** Multiple erythrocyte infection (MEI) as a percentage of overall invasions relative to orbital shaking speed for NF54, 3D7 and Dd2. Shaking speed normalized to clustering threshold for the 6-well plates.

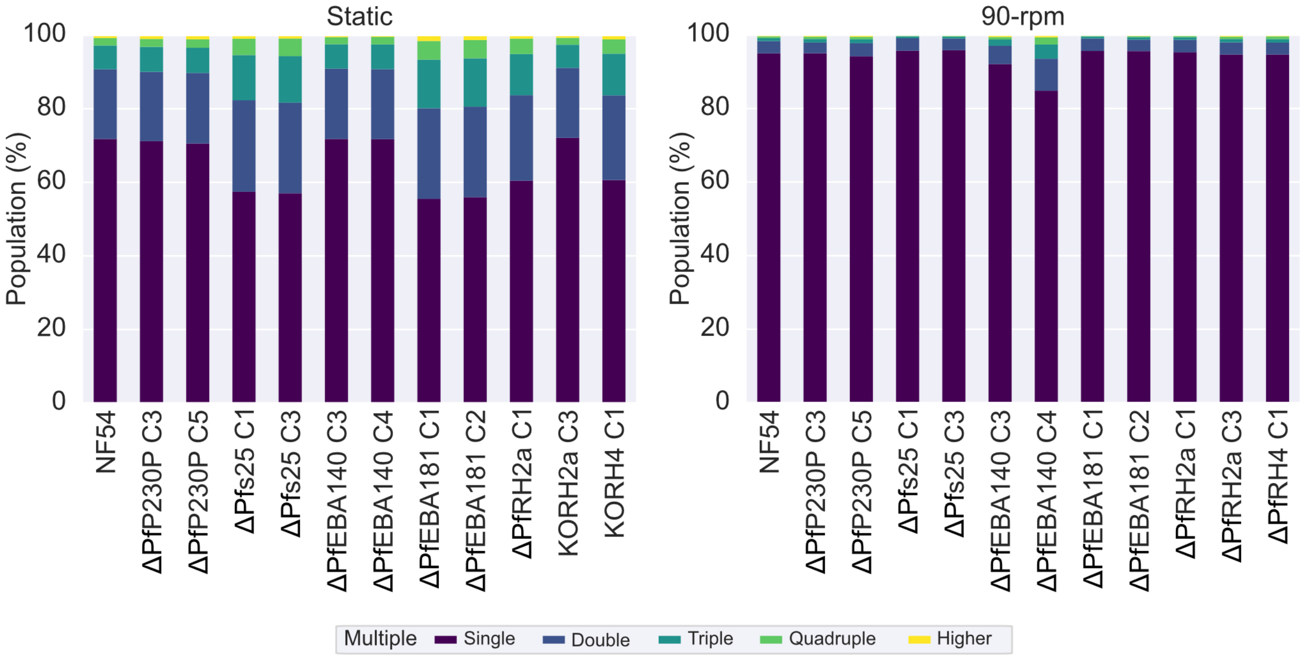


**Fig. S7. Investigating the rate of multiple infections of PfEBA and PfRH knockout lines under static and shaking.** The percentage of ring-stage-infected erythrocytes (excluding higher DNA counts defined as late-stage) that are made up of multiple infections. The graph represents an average of the peaks from the invasion cycle in which the MEI peaks were clearly defined. There is no data for △PfEBA175 as the peaks of the MEI were not clear in any of the samples for the assay in which it was tested.

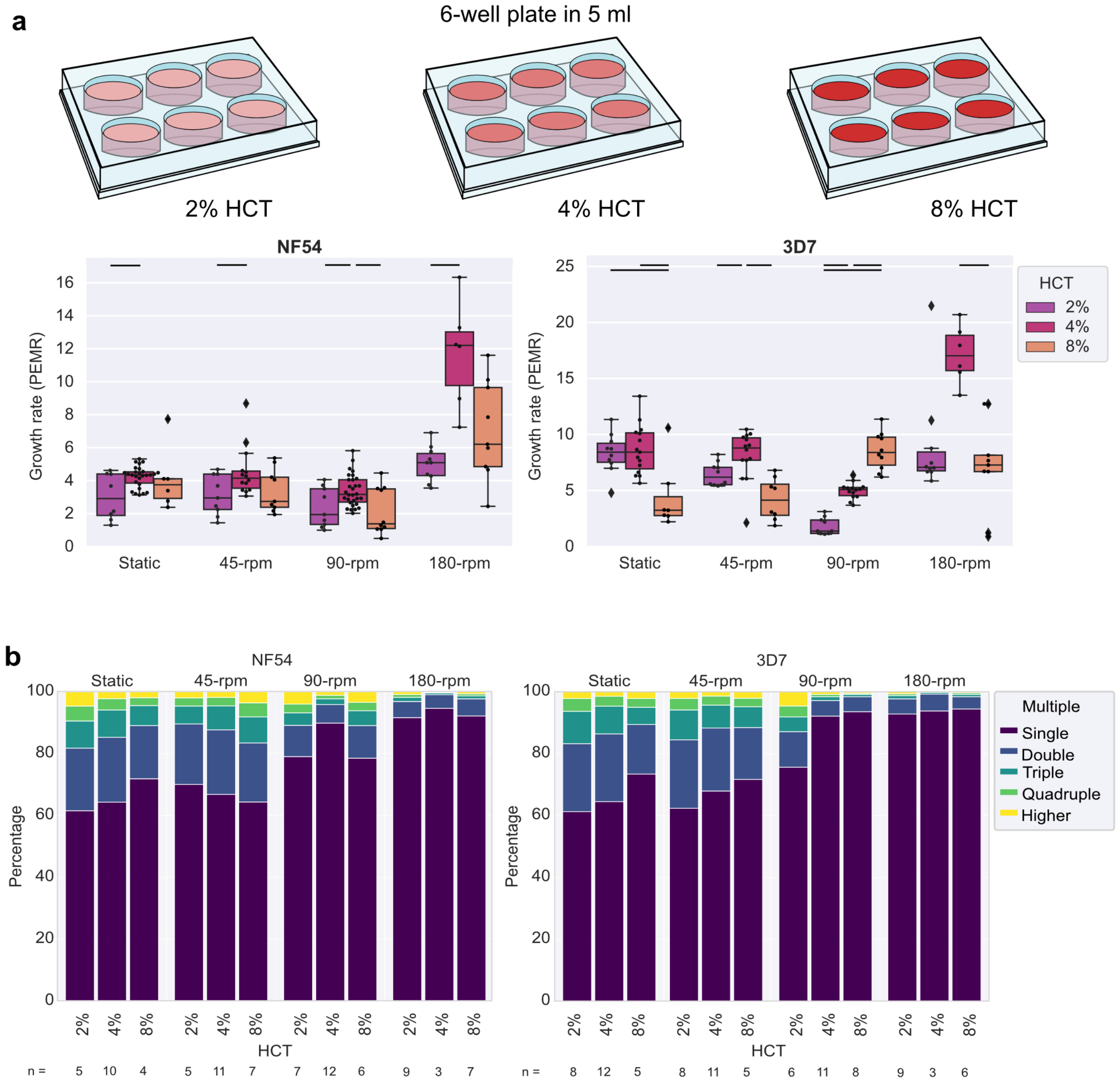


**Fig. S8. Comparison of the effect of different haematocrit on how shaking speed affects the multiple invasion rate and growth of NF54 and 3D7.** All cultures were kept in a 5 ml culture volume in a 6-well plate. All data presented in the figure was collected in parallel. Haematocrit will affect the frequency of contact between merozoites and erythrocytes during the window in which the merozoites are invasion-competent after egress, but it will also affect the density of parasites at a given HCT; a parasitemia of 0.5% will result in twice the density of parasites per culture volume in 4% HCT vs 2% HCT and four times the density in 8% HCT vs 2% HCT. **(a)** Cartoons to represent the different conditions tested. **(b)** The mean growth rate is shown as the parasite erythrocyte multiplication rate (PEMR); each point shows a PEMR measured for a single invasion cycle for an individual well. The lines between conditions indicate the conditions that were significantly different (t-test) at a greater than 5% level of significance. **(c)** Graphs representing the mean frequency of different multiple erythrocyte infections for a given speed for each wild-type line. The frequency of single-infected, double-infected, triple-infected, quadruple-infected, and higher-infected rings is shown as a fraction of the total infected ring-stage parasites. The number of rings per erythrocyte was measured using the peak intensity of the ring-stage infected samples and the number of counts in each peak.

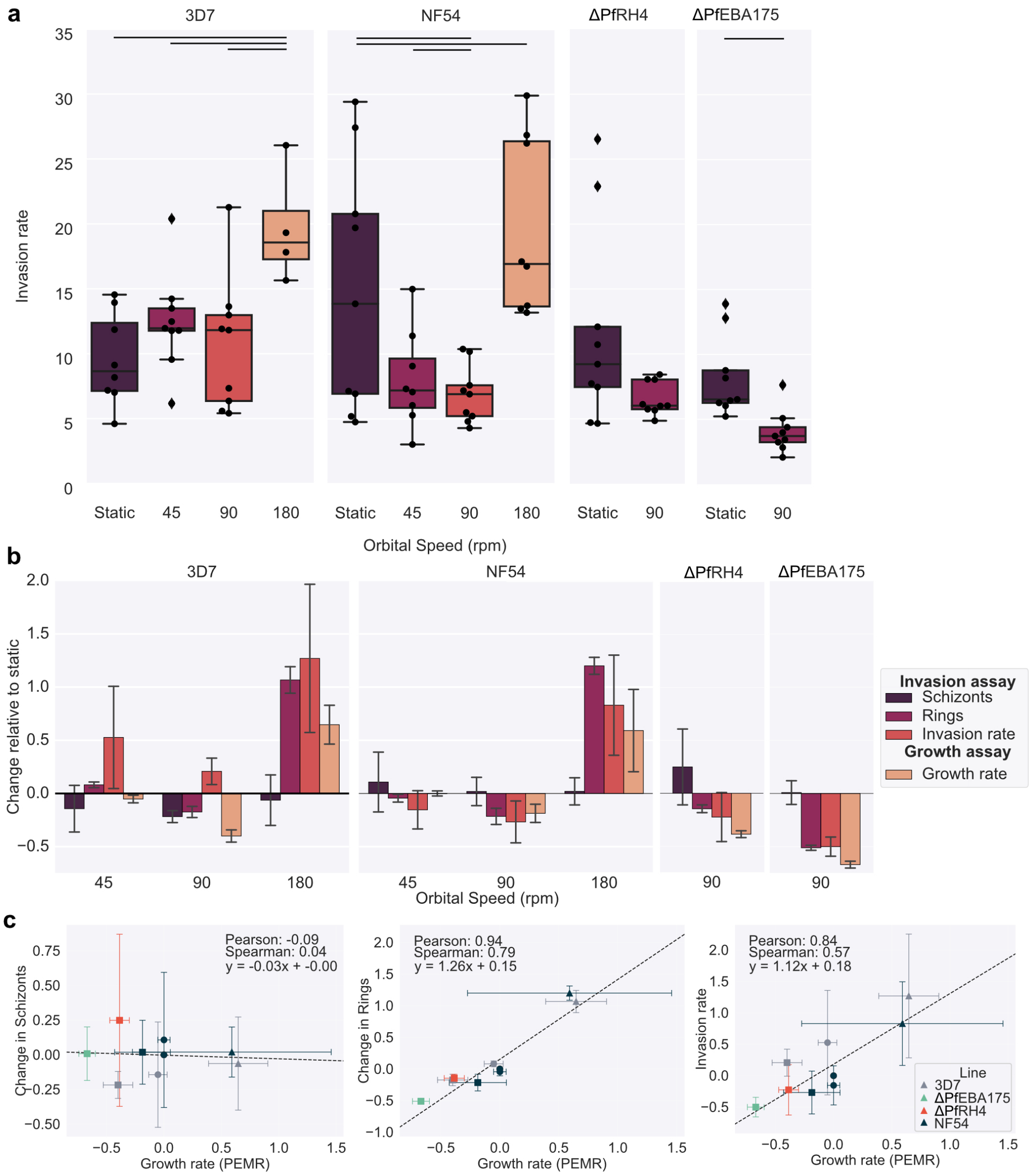


**Fig. S9. Comparing changes in growth due to shaking on effects of shaking on egress and invasion.** All cultures were kept at a 4% HCT in a 5 ml culture in a 6-well plate. All data presented in the figure was collected in parallel. Wild-type lines 3D7 and NF54 were tested along with △PfEBA175c2 and △PfRH4c1, which are in the NF54 background. Measurements were taken before and after a 3.5 hour incubation window. The percentage of rings and schizonts was measured for each well before and after the invasion window. The change in newly invaded RBC rings was calculated by subtracting the percentage of rings present before incubation from the percentage of rings measured after invasion. The change in schizonts was calculated by subtracting the percentage of schizonts present after incubation from the percentage of rings measured before invasion. The invasion rate was calculated by dividing the change in the rings by the change in schizonts. Anything invasion rate of about 35 was excluded as an outlier as there is a maximum of 32 merozoites per schizont. **(a)** Boxplot comparing the invasion rates measured for the different lines across the shaking speed; each point is a single invasion cycle for an individual well. The lines between conditions indicate the conditions that were significantly different (t-test) at a greater than 5% level of significance. **(b)** The change in schizonts, change in rings and invasion rate was compared to the previous measurements of growth rate. All measurements were normalised to the static measurement to allow comparison. **(c)** Scatter plot comparing the normalised growth rates to the normalised change in schizonts, change in rings and invasion rate.

**Movie S1. Top and side view of 6-well plate with 5 ml of media on an orbital shaker as shaking speed is increased. Available at https://zenodo.org/doi/10.5281/zenodo.13372477.**

**Movie S2. Top and side view of 50 ml falsk with 5 ml of media on an orbital shaker as shaking speed is increased. Available at https://zenodo.org/doi/10.5281/zenodo.13372477.**

**Movie S3. Top and side view of 24-well plate with 1 ml of media on an orbital shaker as shaking speed is increased. Available at https://zenodo.org/doi/10.5281/zenodo.13372477.**

**Movie S4. Top and side view of 96-well plate with 100 ul of media on an orbital shaker as shaking speed is increased. Available at https://zenodo.org/doi/10.5281/zenodo.13372477.**
